# Cnidaria XIAP activates caspase-mediated cell death

**DOI:** 10.1101/2024.07.06.602363

**Authors:** Yuan Chen, Meng Wu, Zihao Yuan, Qingyue Wang, Hang Xu, Li Sun

## Abstract

In vertebrate, X-linked inhibitor of apoptosis (XIAP) is a potent inhibitor of apoptosis. XIAP inhibits apoptosis by interacting with proapoptotic caspases via the baculovirus IAP repeat (BIR) domains and mediating caspase ubiquitination via the really interesting new gene (RING) domain and ubiquitin-associated (UBA) domain. In invertebrate, mostly arthropods, XIAP is also known as an apoptosis inhibitor. To date, no study on basal metazoan XIAP has been documented. In the present work, we examined the biological activity of XIAP from jellyfish *Aurelia coerulea* (AcXIAP) and other non-bilaterians. AcXIAP possesses three BIRs and one RING domain but lacks the UBA domain. AcXIAP augmented the apoptosis-inducing activity of all of the four *A. coerulea* caspases, of both the initiator and the effector clades, identified in this study. AcXIAP activated caspase via one of the BIRs, which bound and stabilized the caspase, and the RING domain, which mediated ubiquitination on the p20 subunit of the caspase in a lysine-independent manner. Similar caspase-activating properties were also observed in the XIAP of hydra, coral, and sponge. In hydra, XIAP knockdown markedly decreased cell death induced by an apoptosis inducer. Together these results revealed the unconventional function and working mechanism of XIAP in Cnidaria, and shed new light into the functional and structural evolution of XIAP.

## Introduction

Apoptosis, a type of programmed cell death (PCD), is a fundamental process required for the maintenance of homeostasis under both physiological and pathological conditions. The operation of apoptosis relies on a class of cysteine-dependent aspartic-specific proteases called caspases [1]. In humans and mice, a dozen caspases have been identified. These caspases are divided into two groups, i.e., apoptotic caspases, which are further divided into apoptosis initiator caspases (-2, -8, -9, 10) and effector caspases (-3, -6, -7), and inflammatory caspases (-1, -4, -5, -11, 12), which are involved in inflammatory responses [2]. Caspases are synthesized as inactive zymogens consisting of a pro-domain and a protease domain (CASc), the latter comprising a large subunit (p20) and a small subunit (p10). The pro-domains of effector caspases generally exhibit no structure, while the pro-domains of inflammatory and initiator caspases typically contain either the caspase recruitment domain (CARD) domain or the death effector domain (DED) [1]. During the process of apoptosis, active effector caspases, induced by the mitochondrial or death receptor-mediated cascade signaling, cleave various substrates, including DNA fragmentation factor 45, gelsolin, and poly (ADP-ribose) polymerase, cumulatively leading to apoptosis [3, 4]. Unlike other common regulatory enzymes, such as phosphorylase, methylase, and acetylase [5–7], caspases hydrolyze their substrates directly and hence exert an irreversible effect on the substrates [4]. As a result, the activities of caspases are tightly controlled, either directly through binding to regulatory proteins, or indirectly through transcription or translation [8].

The inhibitor of apoptosis protein (IAP) represents a family of negative regulators of caspase and apoptosis [9]. All IAPs contain at least one copy of the baculovirus IAP repeat (BIR) domain in their N-terminus. Most IAPs contain three BIR domains, which confer on IAPs the ability to bind caspases and other substrates [8]. X-linked inhibitor of apoptosis (XIAP), the most potent and multifaceted member of IAPs, also contains a Ub-associated domain (UBA) for interacting with poly-Ub chains, and a RING (Really Interesting New Gene) domain pivotal for E3 ligase activity [8]. In humans, XIAP inhibits both the effector caspase-3/7 and the initiator caspase-9 [10]. The inhibitory effect of XIAP on caspase-3/7 depends on the binding of the second BIR (BIR2) to the caspase, while the inhibitory effect of XIAP on caspase-9 is mediated by the third BIR (BIR3). The RING domain of XIAP mediates the transfer of the ubiquitin moiety from the enzyme E2 to the substrate caspase, whereby targeting the caspase for proteasomal degradation [8, 9].

Caspases are conserved from invertebrate to vertebrate, and caspase genes have been identified in the genomes of sponge, cnidarians, and all bilaterian clades [11–14]. The functions of caspases have been studied in mammals and the major invertebrate model organisms, the nematode *Caenorhabditis elegans* and the arthropod *Drosophila melanogaster*[13]. In addition, two recent reports showed that coral and abalone caspase-3 activated pyroptosis, a type of PCD different from that of apoptosis [15, 16]. In contrast, functional studies of IAPs in invertebrates are documented mainly with arthropods [17–20]. In insects, as in mammals, IAPs exert an inhibitory effect on caspase-mediated apoptosis [21–23]. In addition to insects, two reports showed that IAPs from other members of Arthropod, i.e., tiger shrimp *Penaeus monodonthe* and Chinese mitten crab *Eriocheir sinensis*, displayed anti-apoptotic activity [24, 25]. These evidences indicate a highly conserved role of IAPs as a negative regulator of apoptosis in both vertebrate and invertebrate.

Cnidaria (corals, sea anemones, jellyfish, and hydroids) is an ancient phylum and a sister group to Bilateria, the two having separated from their common ancestor over 500 million years ago [26]. Genome sequence analyses revealed complex caspase family members in cnidarians [27–29]. However, the functions of the caspases and their regulatory IAPs in Cnidaria remain to be explored. Particularly, in jellyfish, no research on caspase or IAP has been documented. In the present study, we identified four caspases and one XIAP ortholog (AcXIAP) from the moon jellyfish *Aurelia coerulea*. We found that the four jellyfish caspases represented different phylogenetic clades but all exhibited apoptotic activities, which were significantly enhanced, rather than inhibited, by AcXIAP. Similar apoptosis-promoting effects were subsequently discovered in the XIAP homologues of other Cnidaria species and Porifera. We further investigated the working mechanism of AcXIAP, and examined the *in vivo* effect of XIAP knockdown on cell death. Our results uncovered the existence and the functioning mechanism of pro-apoptotic XIAP in basal metazoans.

## Results

### Jellyfish XIAP enhances caspases-mediated apoptosis

Four caspases were identified from jellyfish *A. coerulea* (S1A Fig). Two of these caspases exhibit no apparent structure in the pro-domain and were named AcCaspA and AcCaspB, respectively. The other two caspases possess a CARD domain besides the CASc domain and were designated AcCCaspA (the first capital “C” indicates “CARD”) and AcCCaspB, respectively. Phylogenetic analysis showed that AcCaspA and AcCCaspA were relatively closely related to human/mouse effector caspases (CASP3/7/6), while AcCaspB and AcCCaspB had closer relationships with human/mouse initiator caspases (S1B Fig). AcCaspA, AcCaspB, AcCCaspA, and AcCCaspB contain 315, 462, 437, and 624 amino acid residues, respectively, and share 23.08% identities with each other. All four caspases possess the conserved catalytic motifs of “SHG” and “QACRG” in p20 and the substrate binding “GSWF/Y” motif in p10 (S1A Fig). When expressed in HEK293T cells, all of the four caspases induced marked apoptosis (Fig 1A, S2 Fig). In addition to the caspases, a XIAP structural homologue, named AcXIAP, was also identified in jellyfish (Fig 1B, S3 Fig). AcXIAP contains 779 amino acid residues and forms three conserved BIR domains and one RING domain. Compared with human XIAP (HsXIAP), AcXIAP is 282 amino acids longer, due mostly to the elongated linker regions, and lacks an apparent UBA domain. The predicted three-dimensional (3D) structure of AcXIAP differed considerably from HsXIAP in the C-terminal RING-containing region and in the linkers connecting the BIRs and the RING domains (Fig 1B). To investigate whether AcXIAP, like mammalian XIAP, inhibited caspase activity, ectopic co-expression of AcXIAP and each of the four jellyfish caspases was conducted. Compared with HsXIAP, which markedly inhibited human caspase-9 (HsCasp9)-mediated apoptosis (S4 Fig), AcXIAP significantly increased, rather than decreased, the apoptosis activities of all four jellyfish caspases (Figs 1C-F).

**Fig 1.**
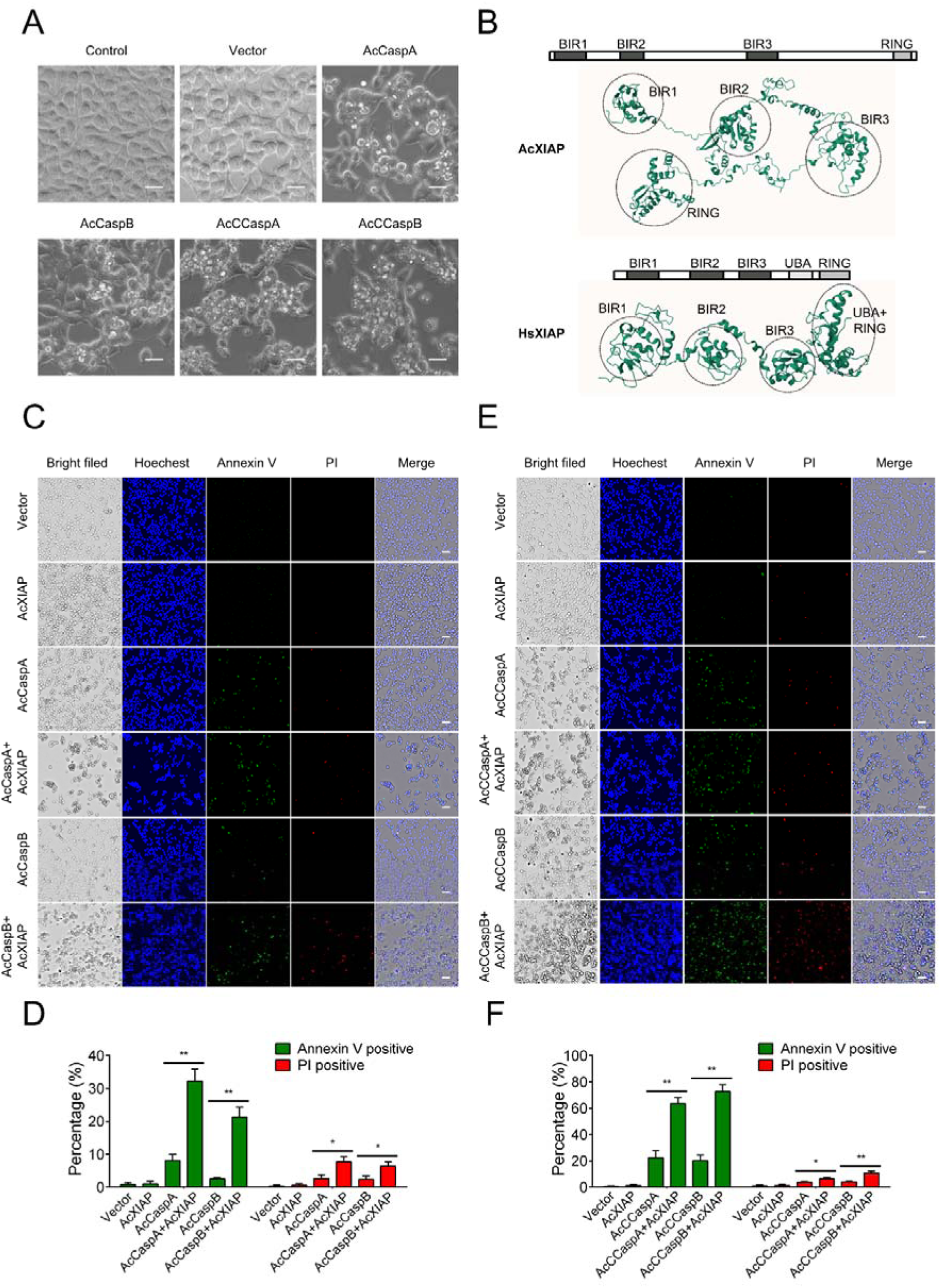
The effect of AcXIAP on caspase-mediated apoptosis. (**A**) HEK293T cells were transfected with or without (control) the backbone vector, or the vector expressing AcCaspA, AcCaspB, AcCCaspA, or AcCCaspB for 30 h, and then subjected to microscopy. Scale bars, 20 μm. (**B**) Schematic representation and predicted structures of AcXIAP and HsXIAP. The conserved domains are circled. (**C-F**) For panels C and D, HEK293T cells were transfected with the backbone vector or the vector expressing AcXIAP, AcCaspA, or AcCaspB, or co-transfected with the vectors expressing AcXIAP and AcCaspA or AcXIAP and AcCaspB for 26 h. For panels E and F, HEK293T cells were transfected with the backbone vector or the vector expressing AcXIAP, AcCCaspA, or AcCCaspB, or co-transfected with the vectors expressing AcXIAP plus AcCCaspA or AcXIAP plus AcCCaspB for 16 h. For panels C-F, the cells were stained with Hoechest 33342, Annexin V-FITC, or PI and then subjected to microscopy (C and E) and analysis of Annexin V/PI-positive cell ratio (D and F). Scale bar, 50 μm. Values are the means of triplicate experiments and shown as means ± SD. \*\**p* < 0.01; \**p* < 0.05.

### AcXIAP promotes caspase activation via BIR3 as well as RING

AcCaspA, the simplest of the four jellyfish caspases in structure, was used to explore the molecular mechanism underlying the caspase-activation effect of AcXIAP. AcXIAP was found to interact and co-localize with AcCaspA (Fig 2A, S5 Fig). Furthermore, in the presence of AcXIAP, both the amount and the activation cleavage of AcCaspA were markedly increased (Fig 2A, panel 4). To determine the specific domain(s) involved in AcXIAP-AcCaspA interaction, a series of AcXIAP truncates were created, i.e., BIR1, BIR2, BIR3, and AcXIAP without the RING domain (AcXIAP-ΔRING) (Fig 2B). The results of co-expression analysis showed that like the wild type AcXIAP, the truncates AcXIAP-ΔRING and BIR3 promoted the activation cleavage of AcCaspA (Fig 2C, panel 3), which was consistent with the Co-IP result showing that AcXIAP-ΔRING and BIR3 were able to interact with and enhanced the activation of AcCaspA (Fig 2C, panel 1). In contrast, BIR1 and BIR2 could not interact with AcCaspA. The activation effect of AcXIAP-ΔRING was apparently dose-dependent (Fig 2D). To further examine the effect of AcXIAP on AcCaspA activity, recombinant AcCaspA was prepared, and its substrate cleavage specificity was determined (S6 Fig). The specific cleavage sequence motif was then served as a substrate to determine the cleavage activity of AcCaspA. The results showed that AcCaspA activity was significantly enhanced by both AcXIAP and AcXIAP-ΔRING; consistently, AcCaspA-mediated apoptosis was significantly enhanced by both AcXIAP and AcXIAP-ΔRING (Figs 2E-G). However, the enhancing effect of AcXIAP-ΔRING was significantly lower than that of AcXIAP, indicating the importance of the RING domain (Figs 2E-G).

**Fig 2.**
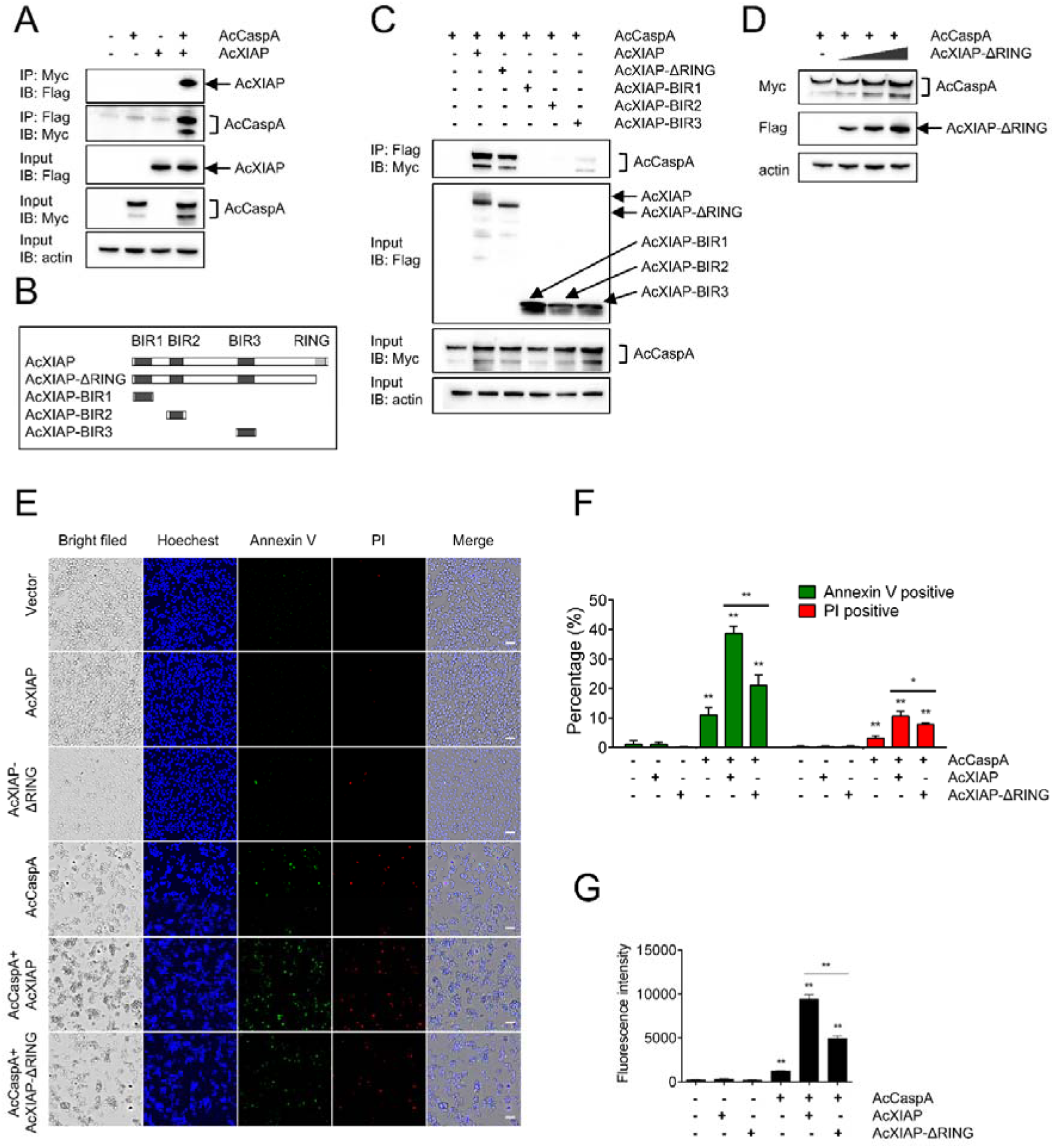
The interaction of AcXIAP with AcCaspA and its effect on AcCaspA activation. (**A**) HEK293T cells were transfected with or without the vector expressing Myc-tagged AcCaspA or Flag-tagged AcXIAP, or co-transfected with the vectors expressing Myc-tagged AcCaspA and Flag-tagged AcXIAP. The cell lysates were immunoprecipitated (IP). The IP and input samples were immunoblotted (IB) with the antibodies against the indicated tags. (**B**) Schematic representation of AcXIAP truncates. (**C**) HEK293T cells expressing Myc-tagged AcCaspA alone or AcCaspA plus various Flag-tagged AcXIAP truncates were immunoblotted as above. (**D**) AcCaspA was expressed alone or together with different doses of AcXIAP-ΔRING in HEK293T cells. The expression levels of AcCaspA and AcXIAP-ΔRING were determined. (**E-G**) HEK293T cells were transfected with or without the vector expressing AcCaspA, AcXIAP, or AcXIAP-ΔRING, or co-transfected with the vectors expressing AcCaspA plus AcXIAP or AcCaspA plus AcXIAP-ΔRING for 26 h. The cells were stained with Hoechest 33342, Annexin V-FITC, PI and then subjected to microscopy (E) and analysis of Annexin V/PI-positive cell ratio (F). The proteolytic activity of AcCaspA was assessed using fluorogenic substrate (G). Scale bar, 50 μm. Values are the means of triplicate experiments and shown as means ± SD. \*\**p* < 0.01; \**p* < 0.05.

### AcXIAP mediates caspase ubiquitination on the p20 subunit

Since in mammalian XIAP, the RING domain functions in ubiquitination, we examined the effect of AcXIAP on AcCaspA ubiquitination. For this purpose, AcCaspA-M, an inactive AcCaspA with mutation at the catalytic site, was created. Co-IP showed that both AcCaspA and AcCaspA-M were ubiquitinated after co-expression with AcXIAP, but not after co-expression with AcXIAP-ΔRING (Figs 3A and B). To determine the AcCaspA region required for ubiquitination, AcXIAP was co-expressed with the p20 or p10 region of AcCaspA. The results showed that p20 was ubiquitinated in the presence of AcXIAP but not in the presence of AcXIAP-ΔRING (Fig 3C). In contrast, AcXIAP or AcXIAP-ΔRING did not affect the ubiquitination of p10, but markedly increased the amount of p10 (Fig 3C). To examine the importance of the lysine residues in AcXIAP-mediated ubiquitination, all of the 28 lysine residues in AcCaspA were mutated. The resulting mutant, AcCaspA-K28, could still be ubiquitinated by AcXIAP (Fig 3D).

**Fig 3.**
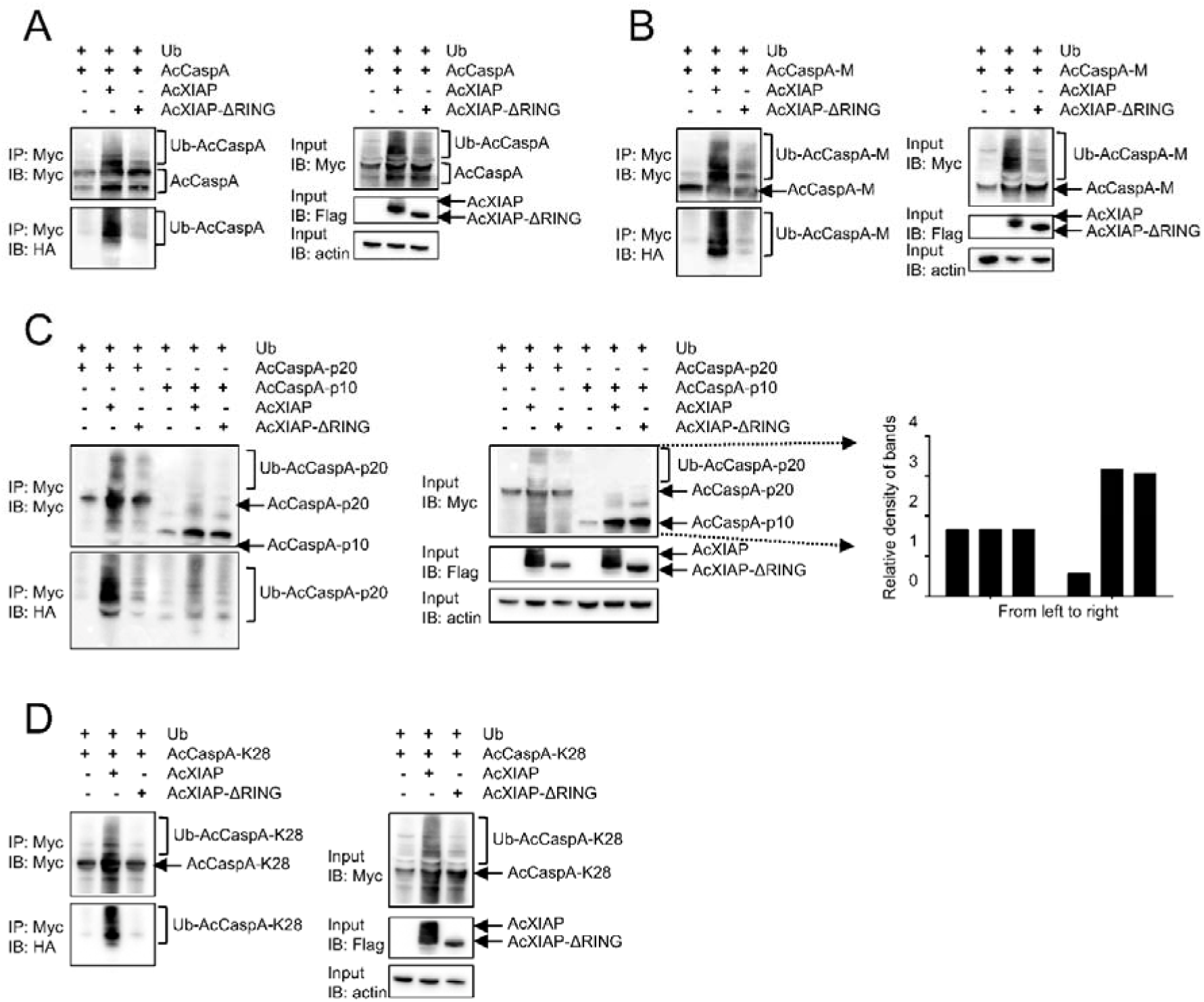
The effects of AcXIAP and AcXIAP-ΔRING on the ubiquitination of AcCaspA variants. For all subfigures, HEK293T cells were transfected with the plasmids expressing the indicated proteins that were tagged as follows: Ub (ubiquitin), HA-tagged; AcCaspA variants, Myc-tagged; AcXIAP and AcXIAP-ΔRING, Flag-tagged. The cell lysates were immunoprecipitated (IP). The IP and input (left and right panels, respectively, in all subfigures) samples were immunoblotted (IB) with the antibodies against the indicated tags. (**A** and **B**) The effects of AcXIAP and AcXIAP-ΔRING on the ubiquitination of AcCaspA (A) and AcCaspA-M (B). (**C**) The effects of AcXIAP and AcXIAP-ΔRING on the ubiquitination of AcCaspA p20 and p10. The relative densities of the p20 and p10 bands in the input samples were calculated with ImageJ. (**D**) The effects of AcXIAP and AcXIAP-ΔRING on the ubiquitination of AcCaspA-K28.

### Activation of caspase-mediated apoptosis is a common feature of non-bilaterian XIAP

Since, as shown above, AcXIAP differed completely from mammalian XIAP in the role of a caspase regulator, we wondered whether this was a general property of basal metazoan XIAP. A search of the available database identified XIAP structural homologs in Porifera (*Amphimedon queenslandica*) and Cnidaria (*Hydra vulgaris* and *Stylophora pistillata*) but not in Ctenophora and Placozoa (Fig 4A, S3 Fig). Like AcXIAP, the Porifera and Cnidaria XIAP homologs possess 2 to 3 BIRs and one RING domain, but lack the UBA domain (S3 Fig). To examine their regulatory effects, the XIAP of sponge, hydra, and coral (named AqXIAP, HvXIAP, and SpXIAP, respectively) were each co-expressed with their respective caspases (AqCasp, HvCasp, and SpCasp, respetively) in HEK293T cells. Subsequent analyses showed that AqXIAP, HvXIAP, and SpXIAP all significantly enhanced caspase-mediated apoptosis (Figs 4B-G, S7 Fig). To examine the effect of XIAP under the *in vivo* condition, hydra polyps with XIAP knockdown were created (S8 Fig). Following treatment with colchicine, an inducer of apoptosis [30], the XIAP-knockdown polyp exhibited much less cell death than the normal polyp (Fig 4H).

**Fig 4.**
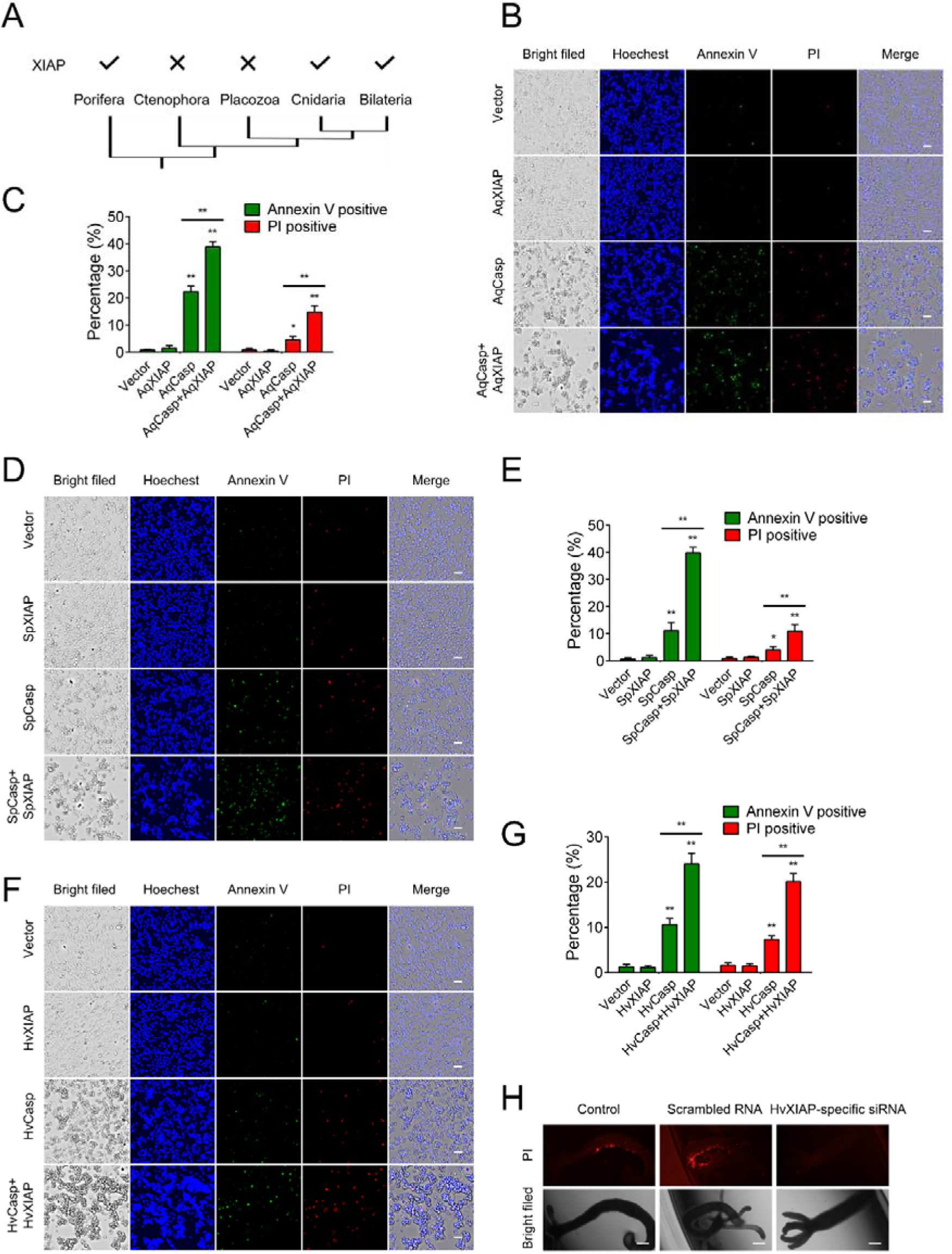
The effects of hydra, coral, and sponge XIAP on caspase-induced apoptosis. (**A**) Metazoan phylogeny with the presence (√) or absence (×) of XIAP ortholog. (**B-G**) HEK293T cells were transfected with the backbone vector or the vector/vectors expressing the indicated sponge caspase/XIAP (B and C), coral caspase/XIAP (D and E), or hydra caspase/XIAP (F and G) for 27 h. The cells were stained with Hoechest 33342, Annexin V-FITC, PI and then subjected to microscopy (B, D, and F) and analysis of Annexin V/PI-positive cell ratio (C, E, and G). Scale bar, 50 μm. Values are the means of triplicate experiments and shown as means ± SD. \*\**p* < 0.01; \**p* < 0.05. (**H**) Hydra polyps were treated with or without (control) scrambled siRNA or HvXIAP-specific siRNA and then stained with propidium iodide (PI). The polyps were observed with a fluorescence microscope. Scale bar, 200 μm.

## Discussion

The moon jellyfish *Aurelia* is an emblematic medusa in Cnidaria, with a complex life cycle like most medusozoans [31]. Over twenty species of *Aurelia* have been recognized [32, 33]. Among them, *A. coerulea* and *Aurelia aurita* are invasive species widely distributed around the Northwest Pacific and Atlantic coast of Europe [33]. In contrast to the invertebrate model organisms *C. elegans* and *Drosophila*, cnidarians (hydra and coral) appear to have complex caspase repertoires [11, 34], but the functions of these caspases remain elusive. In jellyfish, whose highly watery and gelatinous nature renders cellular and *in vivo* studies difficult, no study on caspase or IAP has been documented. In the present study, we identified and characterized four *A. coerulea* caspases, all of which were found to induce robust apoptosis when expressed in human cells, suggesting that these caspases probably worked via conserved mechanisms similar to that of mammalian caspases. This is consistent with the presence of highly conserved catalytic motifs and substrate binding sequences in the four jellyfish caspases. Nevertheless, it should be noted that although AcCCaspA, one of the four identified jellyfish caspases, was grouped with human/mouse effector caspases, it differs from mammalian effector caspases by possessing an N-terminal CARD domain. On the other hand, AcCaspB, which possesses neither CARD nor DED domain, proved to be more closely related to human/mouse initiator caspases than to the effector caspases. These results indicate that the caspase classification criteria of the vertebrate cannot be mechanically applied to invertebrate species.

XIAP is the most studied members of the IAP family and well-known as a potent inhibitor of apoptosis in vertebrates and some invertebrates, mostly arthropods [8, 17, 35]. In this study, we found that AcXIAP significantly enhanced, rather than inhibited, the apoptotic activities of all of the four examined jellyfish caspases. Since the four jellyfish caspases differ considerably in sequence characteristics and fall into different phylogenetic clades, the activating effect of AcXIAP is probably not specific to a certain type of caspase. Similar to human XIAP, AcXIAP contains the conserved RING domain and three BIR domains, however, AcXIAP differs noticeably from human XIAP in the predicted 3D structures over the RING region and the linker regions, especially in the linkers bridging BIR2-BIR3 and BIR3-RING. These linkers, unlike that in human XIAP, are much longer and adopt, in parts, well-defined secondary structures, and connect the BIRs and RING modules loosely into a higher-ordered structure more extended than that of human XIAP. It is reasonable to speculate that these structural “peculiarities” may account for, or at least contribute to, the unconventional quality of AcXIAP.

A previous study showed that the BIR3 domain of human XIAP directly interacted with and inhibited caspase 9 [10]. For AcXIAP, we found that it was also BIR3 that mediated the binding to the target caspase, though in this case the binding leading to a stimulating, rather than inhibiting, effect on caspase activation. The observation that BIR3 alone could reproduce the caspase-activating effect of AcXIAP not only highlighted the vital role of BIR3, but also indicated that BIR3 could play its role in an autonomous manner that did not require the cooperation of the other domains of AcXIAP. Like AcXIAP, BIR3 enabled more AcCaspA, in both uncleaved and cleaved forms, to be detected, suggesting that BIR3 probably facilitated caspase activation by holding the caspase in a more stable state. In addition to BIR3, another AcXIAP truncate, AcXIAP-ΔRING, also exerted an augmenting effect on AcCaspA activation, but this effect was significantly lower than that of AcXIAP. Since the RING domain in mammalian XIAP mediates ubiquitination [36], it is possible that RING-associated ubiquitination is involved in AcXIAP regulation of caspase activation. Indeed, we found that AcXIAP ubiquitinated AcCaspA at p20 in a RING-dependent manner, thus indicating a conserved function of RING in invertebrate and vertebrate XIAP. However, the ubiquitination mechanism of the RING of AcXIAP is probably different from that of vertebrate XIAP, since mutation analyses demonstrated that none of the 28 lysine residues in AcCaspA was required for AcXIAP-mediated ubiquitination.

The IAPs, characterized by at least one BIR domain and a RING motif, occurred early in metazoan evolution [37, 38]. In this study, we found that besides jellyfish, XIAP structural homologs existed in other non-bilaterian organisms, including sponge, hydra, and coral. It is striking that, like AcXIAP, all of these non-bilaterian XIAPs exhibited caspase-activating property, suggesting a conserved pro-apoptosis function of XIAP in basal metazoans. This was supported by the *in vivo* study in hydra, which showed that XIAP knockdown markedly weakened colchicine-induced cell death. With these observations, one may speculate that XIAP in its early forms probably act as a caspase activator via its BIR-mediated substrate binding and RING-mediated ubiquitination, the latter in a lysine-independent manner. It is possible that the BIR may exert, in a chaperon-like manner, on the substrate caspase a stabilizing effect that is further strengthened by subsequent ubiquitination. It is also possible that both BIR interaction and RING-mediated ubiquitination may alter the structure of the bound caspase in a way that directly affects the activation cleavage of the caspase.

In conclusion, in this study we discovered the existence and the working mechanism of pro-apoptosis XIAP in Cnidaria. Our results indicated that the basic modular organization of XIAP had come into existence in non-bilaterians, and the co-operation of the different task-oriented domains enables XIAP to play the role of a broad range caspase activator. In Bilateria, the general domain-based functional division of the primitive XIAP is retained, which preserves the nature of XIAP as a caspase regulator, but evolution-selected structural alterations have led to a role-reversal of the XIAP from a positive regulator to a negative regulator. These findings shed new light on the function, evolution, and regulation mechanism of XIAP.

## Materials and Methods

### Animals and cell lines

Jellyfish *Aurelia coerulea* medusas were maintained in the laboratory as described previously [39]. The entire body of the animal was used for RNA isolation with a TRIZOL reagent (Invitrogen, USA). Hydra polyps were collected from Beijing, China, and cultured at 18°C in Hydra medium composed of 1 mM NaCl, 1 mM CaCl_2_, 0.1 mM KCl, 0.1 mM MgSO_4_, and 1 mM tris-HCl (pH 7.6). The animals were fed with fresh *Artemia nauplii* larvae as previously reported [29]. HEK293T cells (American Type Culture Collection) were cultured in Dulbecco’s modified Eagle’s medium supplemented with 10% fetal bovine serum (Gibco, Renfrewshire, UK) at 37°C in a 5% CO_2_ incubator.

### Sequence and phylogenetic analysis

For sequence alignment, XIAP sequences (S1 Table) were selected from the National Center for Biotechnology Information (NCBI) (www.ncbi.nlm.nih.gov) and aligned using DNAMAN. Phylogenetic analysis was performed as reported previously using IQ-TREE 2 v.2.1.2 with JTT+R3 substitution models and 1000 bootstraps [40]. Conserved domain analysis was performed using the conserved domain database of NCBI (www.ncbi.nlm.nih.gov/Structure/cdd/wrpsb.cgi).

### Gene cloning and mutagenesis

*A. coerulea* caspases and XIAP genes were identified by PCR with primers (S2 Table) based on the caspase and XIAP sequences of *Aurelia aurita*. The PCR products were cloned into pEASY-T1 Simple using the T1 Simple-cloning Kit (TransGen, Beijing, China) and verified by sequencing. The codon-optimized protein coding sequences (CDSs) of sponge, hydra, coral and human caspases/XIAPs (S1 Table) were synthesized by Sangon Biotech (Shanghai, China). The CDSs of AcCaspA and AcXIAP truncates were obtained based on their respective full-length CDSs. Site-directed mutagenesis of lysine to arginine and cysteine to alanine in AcCaspA was performed using the Fast Mutagenesis System (TransGen, Beijing, China) according to the manufacturer’s instruction. For transfection, the CDSs were subcloned into pmCherry-N1, pEGFP-N1, pCAGGS-Flag, or pCS2-6 × Myc expression vector (Clontech, Mountain View, CA, USA). All plasmids were verified by sequencing.

### Detection of cell death

HEK293T cells were seeded in 24-well plates (Costar, Corning, NY, USA) and transfected with indicated caspase and/or XIAPs. At 24-36 h post transfection, the cells were washed with PBS and then stained with Hoechest 33342, Annexin V-FITC and propidium iodide (PI). Image acquisition and analysis were performed with automatic real-time live cell imaging detection system (Spark®Cyto, Tecan, Switzerland).

### Preparation of recombinant protein

The CDS of AcCaspA was subcloned into pET-28a (+) expression vector (Novagen, Madison, WI, USA) and expressed as His-tagged protein. The recombinant plasmid was introduced into *Escherichia coli* Transetta (DE3) (TransGen, Beijing, China) via transformation. The recombinant protein was prepared from the *E. coli* transformant as described previously [41] with Ni-NTA agarose (QIAGEN, Dusseldorf, Germany). The protein purity was determined by SDS-PAGE (S6 Fig).

### AcCaspA activity

To determine the proteolytic activity of AcCaspA, AcCaspA was incubated with various colorimetric substrates as described previously [42]. After 1.5 h incubation, the release of ρ-nitroanilide (ρNA) was monitored at OD_405_ nm. To determine the AcCaspA activity in AcCaspA-expressing HEK293T cells, the cell lysates were buffered in a 100 μl reaction system [50 mM Hepes (pH 7.5), 3 mM EDTA, 150 mM NaCl, 0.005% (v/v) Tween 20, and 10 mM dithiothreitol (DTT)] and incubated with Ac-VDVAD-AFC (MedChem Express, NJ, USA) at a final concentration of 200 μM at 25°C for 1.5 h. Fluorescence was then measured with a BioTek Synergy HT plate reader (BioTek Instruments, VT, USA).

### XIAP**−**caspase interaction

HEK293T cells were seeded in 24-well plates (Costar, Corning, NY, USA) overnight. The cells were transfected with the plasmid expressing caspases, XIAP, or caspase plus XIAP using PolyJet transfection reagent (SignaGen Laboratories, Ijamsville, MD, USA). After various hours of transfection, the cells were observed with an inverted microscope (TiS/L100, Nikon, Tokyo, Japan). For co-localization analysis, HEK293T cells were co-transfected with the vectors expressing EGFP-tagged AcCaspA and mCherry-tagged AcXIAP for 24 h, and the cells were observed with a Carl Zeiss LSM 710 confocal microscope (Carl Zeiss, Jena, Germany).

### HvXIAP knockdown and its effect on cell death

HvXIAP-specific small interfering RNA (siRNA) (5’-GGACCAUAAUCCAAUUCAATT-3’) and scrambled RNA (5’-UUCUCCGAACGUGUCACGUTT-3’) (negative control) were designed and synthesized by Sangon Biotech (Shanghai, China). Hydra polys were randomly divided into different groups (10 animals/group). The groups were administered with or without HvXIAP-specific siRNA or scrambled RNA with an electroporator (BTX Gemini X2, USA) based on the reported method of electroporation [43]. After electroporation, the polyps were transferred into Hydra medium supplemented with 20% (v/v) hyperosmotic dissociation medium consisting of 3.6 mM KCl, 6 mM CaCl_2_, 1.2 mM MgSO_4_, 6 mM sodium pyruvate, 6 mM sodium citrate, 12.5 mM N-tris [hydroxymethyl] methyl-2aminoethanesulfonic acid (TES), 6 mM glucose, and 50 mg/mL rifampicin. After 24 h incubation, the medium was replaced by fresh Hydra medium. To examine the efficiency of electroporation, hydra polyps were electroporated as above with or without 6-carboxy-fluorescein-labeled scrambled RNA and observed with a fluorescence microscope (Olympus, Tokyo, Japan) at 24 h after electroporation. For HvXIAP knockdown, hydra polyps were electroporated as above with HvXIAP-specific siRNA or scrambled RNA. The knockdown efficiency was determined by Quantitative Real-Time Reverse Transcription-PCR (qRT-PCR) as reported previously [44] with primers listed in S2 Table. To examine the effect of HvXIAP knockdown on cell death induced by colchicine, hydra polyps were administered with HvXIAP-specific siRNA or scrambled RNA as above and then treated with colchicine (2 mg/ml) for 2 h, followed by washing in Hydra medium. The polyps were stained with PI (2 μg/ml) for 15 min and then observed with a fluorescence microscope (Olympus, Tokyo, Japan).

### Immunoblotting

HEK293T cells transfected with indicated plasmids were lysed with lysis buffer (Beyotime, Shanghai, China), and the lysate was boiled in SDS loading buffer and electrophoresed in 12% SDS-PAGE (GenScript, Piscataway, NJ). The proteins were transferred to a NC membrane and treated with the indicated antibodies. β-actin served as a control. The protein bands were visualized with an ECL kit (Sparkjade Biotechnology Co. Ltd, Shandong, China). The antibodies used for immunoblotting are as follows: HRP-conjugated mouse anti DDDDK-Tag mAb, HRP-conjugated mouse anti Myc-Tag mAb, HRP-conjugated mouse anti HA-Tag mAb, and HRP-conjugated β-actin mouse mAb. All antibodies were from ABclonal Technology, Wuhan, China.

### Co-immunoprecipitation (Co-IP)

For Co-IP, HEK293T cells were transfected with the indicated plasmids for 24 h and lysed with lysis buffer. The cells lysates were incubated with anti-Flag or anti-Myc affinity gels (Beyotime, Shanghai, China) 4°C overnight. The affinity gels were then washed with PBS for 10 times and boiled with loading buffer to elute the bound proteins. The eluted proteins were subjected to immunoblotting as described above.

### Statistical analysis

The GraphPad Prism software 6 was used for statistical analysis, with statistical significance defined as *p* < 0.05. The unpaired Student’s t-test was used for the comparison between two groups. Error bars represent the mean ± SD.

## Supporting information

**S1 Fig. Phylogenetic analysis of jellyfish caspases.** (A) Schematic representation of the four jellyfish caspases. (B) Phylogenetic analysis of jellyfish and human/mouse caspases. Branch support values are expressed as percentages.

**S2 Fig. Apoptotic activity of jellyfish caspases. (A** and **B)** HEK293T cells were transfected with or without (control) the backbone vector, or the vector expressing AcCaspA or AcCaspB for 28 h. (**C** and **D**) HEK293T cells were transfected with or without (control) the backbone vector, or the vector expressing AcCCaspA or AcCCaspB for 16 h. For panels A-D, the cells were stained with Hoechest 33342, Annexin V-FITC, or PI and then subjected to microscopy (A and C) and analysis of Annexin V/PI-positive cell ratio (B and D). Scale bars, 50 μm. Values are the means of triplicate experiments and shown as means ± SD. \*\**p* < 0.01.

**S3 Fig. Comparative sequence analysis of non-bilaterian XIAP and human XIAP (HsXIAP).** (**A**) Schematic representation of non-bilaterian XIAP and HsXIAP. (**B**) Alignment of non-bilaterian XIAP and HsXIAP. The identical residues are indicated in black; the residues that are ≥75% and ≥50% identical are indicated in red and blue, respectively. The black boxes indicate the conserved BIR and RING domains.

**S4 Fig. The effect of HsXIAP on HsCasp9-mediated apoptosis.** HEK293T cells were transfected with the backbone vector or the vector expressing HsCasp9 or HsXIAP, or co-transfected with the vectors expressing HsCasp9 and HsXIAP for 26 h. The cells were stained with Hoechest 33342, Annexin V-FITC, or PI and then subjected to microscopy (A) and analysis of Annexin V/PI-positive cell ratio (B). Scale bar, 50 μm. Values are the means of triplicate experiments and shown as means ± SD. \*\**p* < 0.01.

**S5 Fig. Co-localization of AcXIAP with AcCaspA.** HEK293T cells were co-transfected with the vectors expressing EGFP-tagged AcCaspA and mCherry-tagged AcXIAP. The cells were observed with a confocal microscope. Scale bar, 5 μm.

**S6 Fig. Analysis of the proteolytic specificity of AcCaspA.** (**A**) Recombinant AcCaspA was analyzed by SDS-PAGE and stained with CBB R-250. M, marker. (**B**) AcCaspA was incubated with different colorimetric caspase substrates, and the released ρNA was measured at 405 nm. Values are the means of triplicate experiments and shown as means ± SD.

**S7 Fig. The effects of hydra, sponge and coral XIAP on caspase-induced apoptosis.** HEK293T cells were transfected with the backbone vector or the vector/vectors expressing the indicated hydra caspase/XIAP (A), sponge caspase/XIAP (B), or coral caspase/XIAP (C) for 27 h. The cells were then subjected to microscopy. Scale bars, 20 μm.

**S8 Fig. Analysis of the efficiency of HvXIAP knockdown.** (**A**) Hydra polyps were electroporated with or without (control) FAM-labeled siRNA and observed with an inverted fluorescence microscope. Scale bar, 200 μm. (**B**) Hydra polyps were electroporated with or without (control) HvXIAP-specific siRNA or scrambled RNA. The expression of HvXIAP was determined by qRT-PCR at 24 h post electroporation. Values are the means of triplicate experiments and shown as means ± SD. \**p* < 0.05.

**S1 Table. List of protein accession numbers.**

**S2 Table. List of primers used for gene cloning.**

## Acknowledgements

This work was supported by the Science & Technology Innovation Project of Laoshan Laboratory (LSKJ202203000) and the Innovation Research Group Project of the National Natural Science Foundation of China (42221005). The authors acknowledge the Oceanographic Data Center, IOCAS, and System Simulation of Laoshan Laboratory for providing the computing resources.

## Author Contributions

Conceptualization and Funding acquisition: LS; Data curation: YC and HX; Methodology: YC; Validation: YC and HX; Investigation: YC, MW, ZHY, and QYW; Formal analysis: YC; Writing - original draft: YC; Writing - review & editing: LS and HX.

## Data Availability

The data presented in this study can be accessed in the article/Supplemental Materials.

