## Supplemental materials for "Cnidaria XIAP activates caspase-mediated cell death"

**S1 Fig**

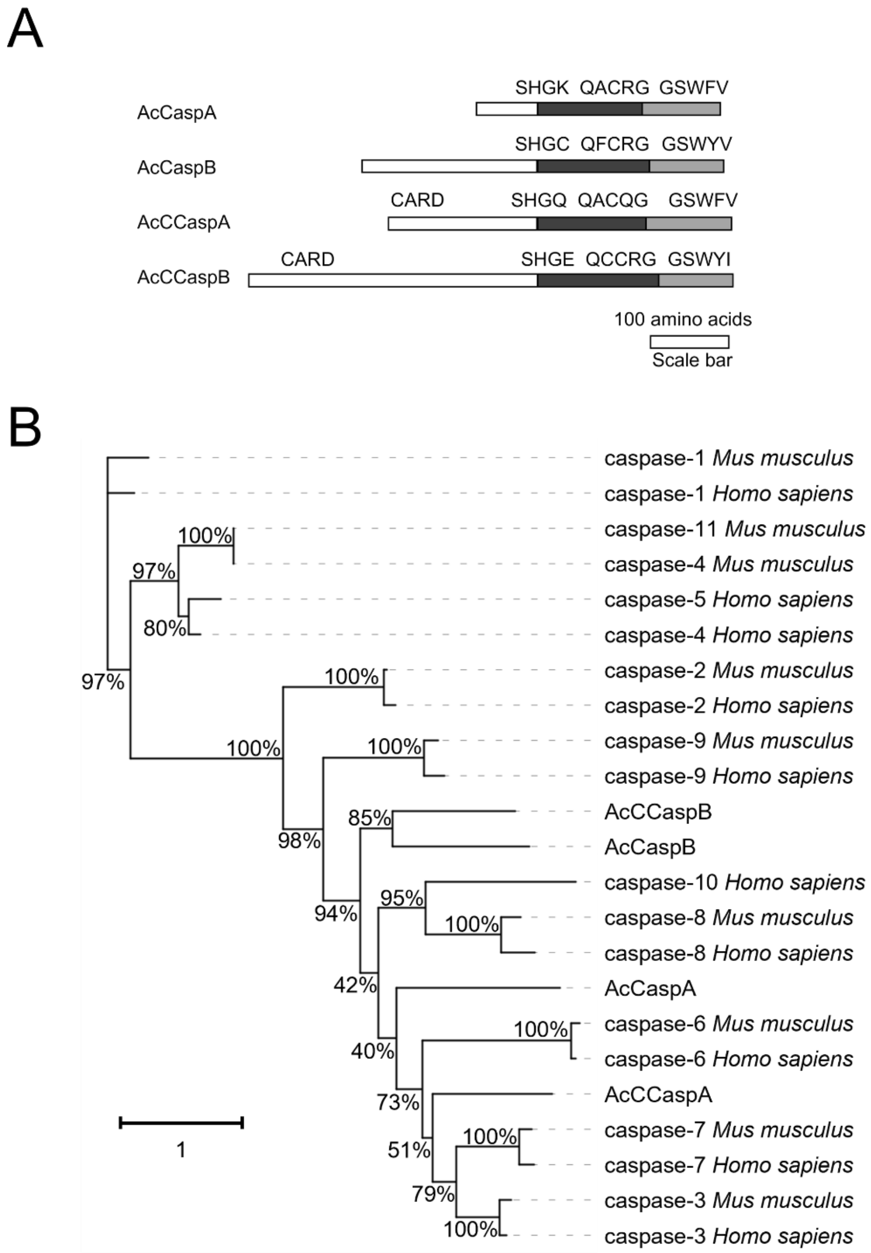
A
SHGK QACRG GSWFV
AcCaspA
SHGC QFCRG GSWYV
AcCaspB
CARD
SHGQ QACQG GSWFV
AcCCaspA
CARD
SHGE QCCRG GSWYI
AcCCaspB
100 amino acids
Scale bar
B
caspase-1 Mus musculus
caspase-1 Homo sapiens
caspase-11 Mus musculus
caspase-4 Mus musculus
caspase-5 Homo sapiens
caspase-4 Homo sapiens
caspase-2 Mus musculus
caspase-2 Homo sapiens
caspase-9 Mus musculus
caspase-9 Homo sapiens
AcCCaspB
AcCaspB
caspase-10 Homo sapiens
caspase-8 Mus musculus
caspase-8 Homo sapiens
AcCaspA
caspase-6 Mus musculus
caspase-6 Homo sapiens
AcCCaspA
caspase-7 Mus musculus
caspase-7 Homo sapiens
caspase-3 Mus musculus
caspase-3 Homo sapiens
100%
97%
80%
97%
100%
100%
100%
98%
85%
94%
95%
100%
42%
100%
40%
73%
100%
51%
79%
100%
1

**S2 Fig**

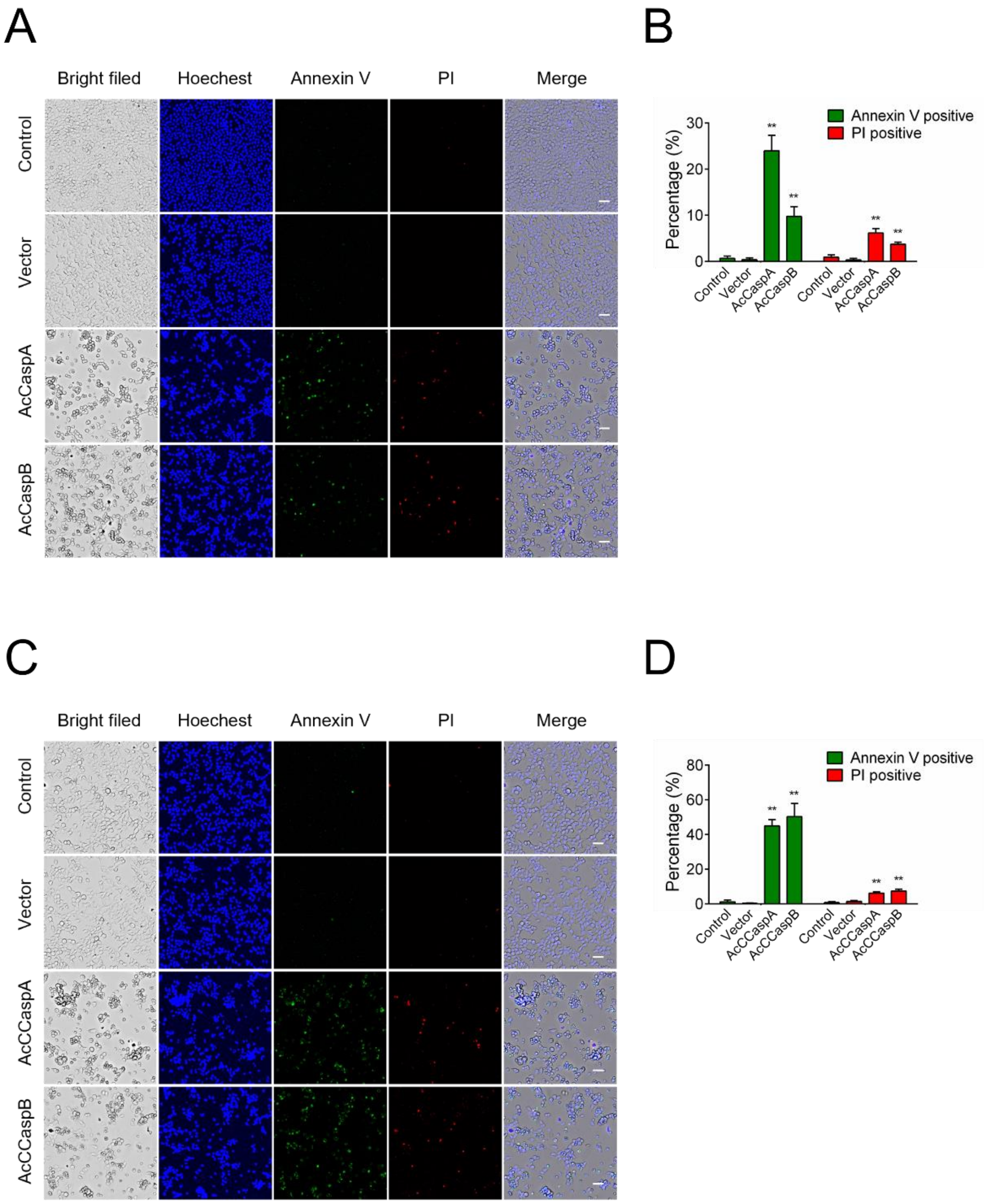
A
Bright filed
Hoechest
Annexin V
PI
Merge
Control
Vector
AcCaspA
AcCaspB
B
Annexin V positive
PI positive
Percentage (%)
30
20
10
0
**
Control
Vector
AcCaspA
AcCaspB
C
Bright filed
Hoechest
Annexin V
PI
Merge
Control
Vector
AcCCaspA
AcCCaspB
D
Annexin V positive
PI positive
Percentage (%)
80
60
40
20
0
**
Control
Vector
AcCCaspA
AcCCaspB

**S3 Fig**

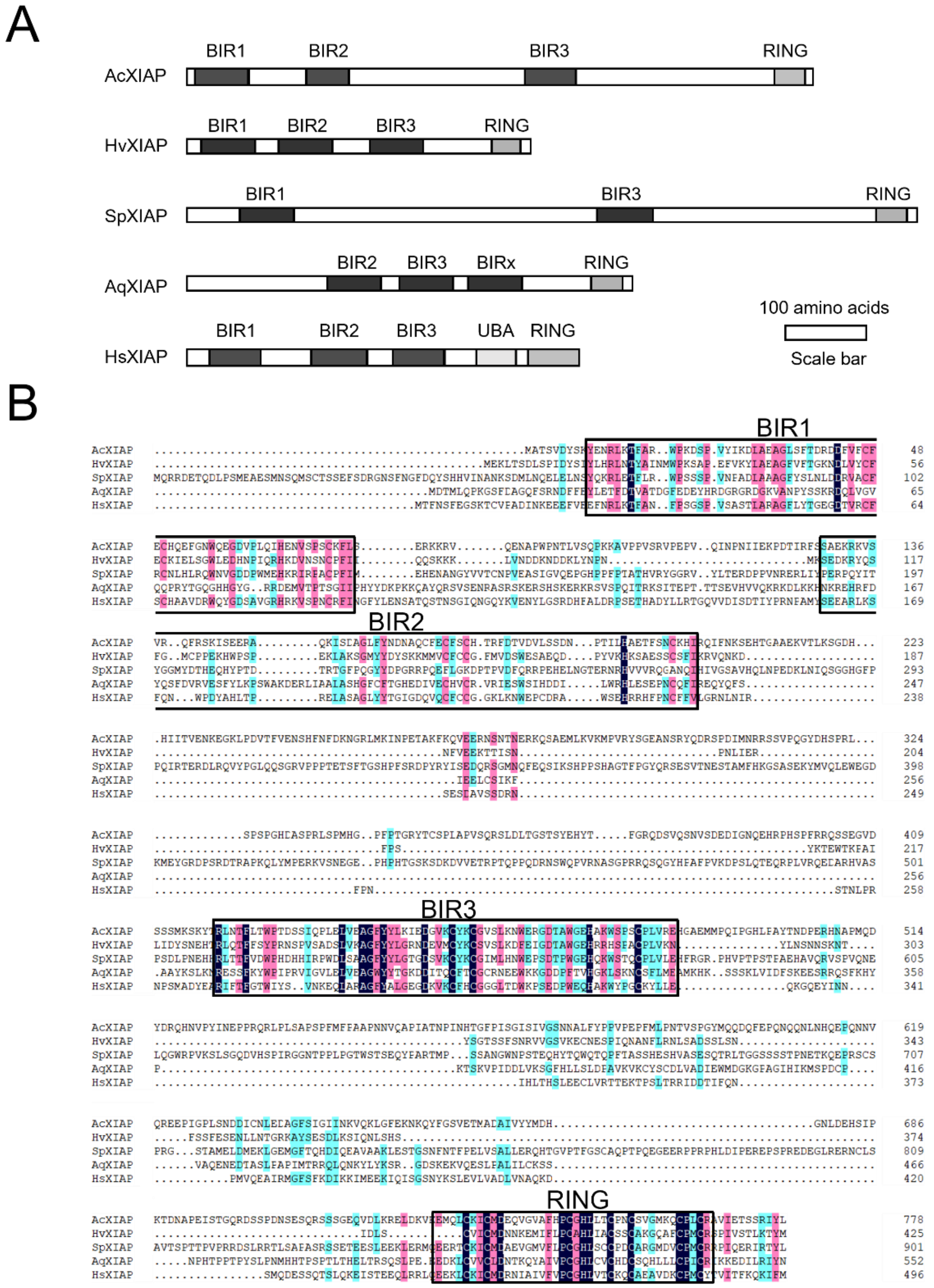
A
BIR1
BIR2
BIR3
RING
AcXIAP
HvXIAP
SpXIAP
BIRx
AqXIAP
UBA
HsXIAP
100 amino acids
Scale bar
B
BIR1
BIR2
BIR3
RING

**S4 Fig**

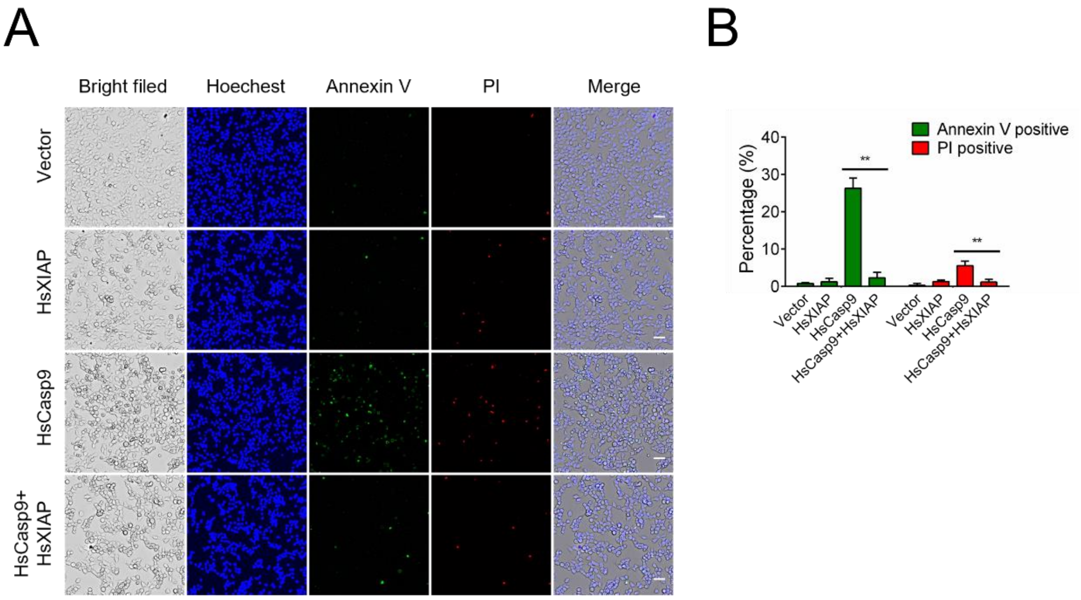
A
Bright filed
Hoechest
Annexin V
PI
Merge
Vector
HsXIAP
HsCasp9
HsCasp9+
HsXIAP
B
Annexin V positive
PI positive
Percentage (%)
40
30
20
10
0
**
**
Vector
HsXIAP
HsCasp9
HsCasp9+HsXIAP
Vector
HsXIAP
HsCasp9
HsCasp9+HsXIAP

**S5 Fig**

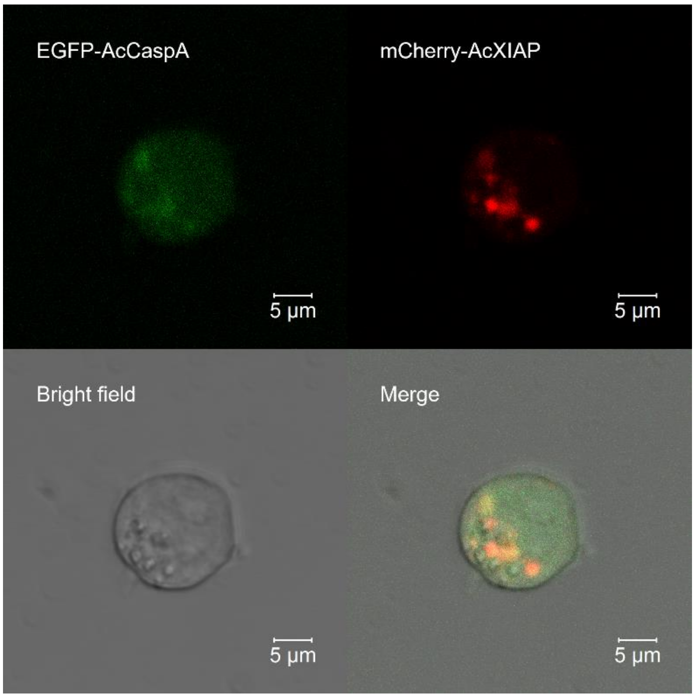
EGFP-AcCaspA
mCherry-AcXIAP
Bright field
Merge
5 μm
5 μm
5 μm
5 μm

**S6 Fig**

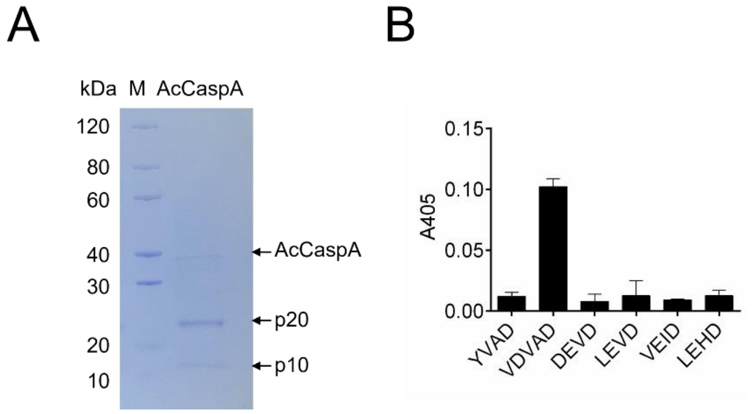
A
kDa
M
AcCaspA
120
80
60
40
30
20
10
AcCaspA
p20
p10
B
0.15
0.10
0.05
0.00
A405
YVAD
VDVAD
DEVD
LEVD
VEID
LEHD

**S7 Fig**

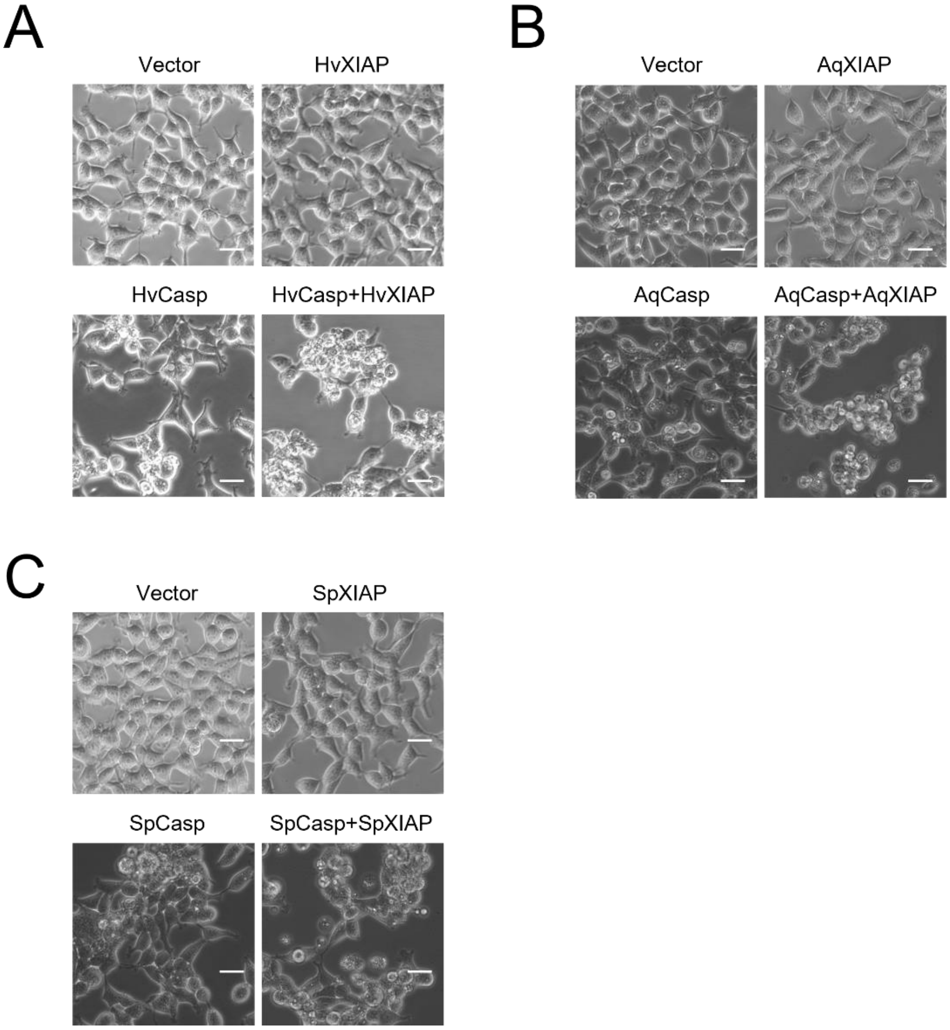
A
Vector
HvXIAP
HvCasp
HvCasp+HvXIAP
B
Vector
AqXIAP
AqCasp
AqCasp+AqXIAP
C
Vector
SpXIAP
SpCasp
SpCasp+SpXIAP

**S8 Fig**

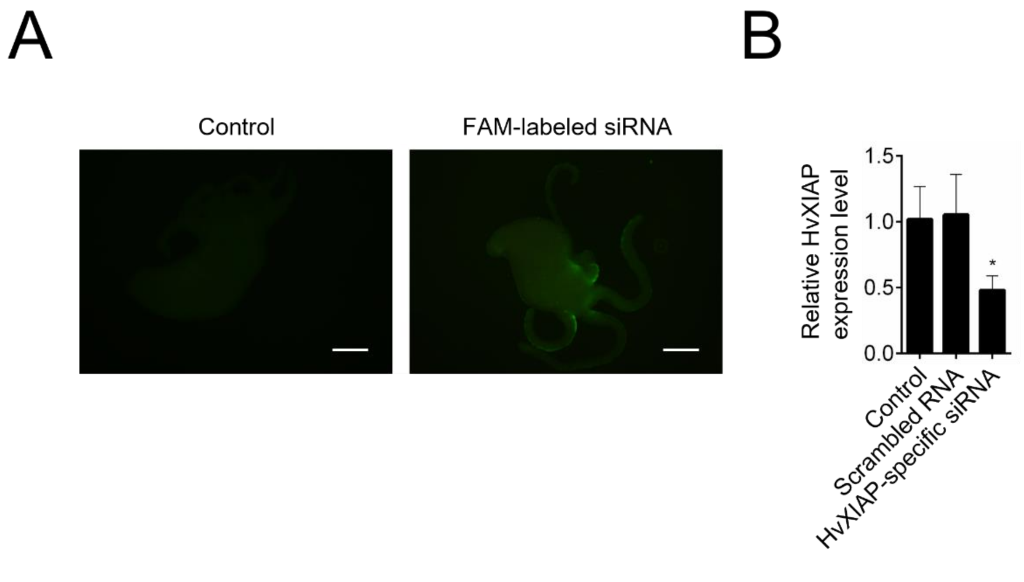
A
B
Control
FAM-labeled siRNA
Relative HvXIAP expression level
1.5
1.0
0.5
0.0
*
Control
Scrambled RNA
HvXIAP-specific siRNA

**S1 Table**

| Protein | Accession |
|---|---|
| HvCasp | >NP_001267785.1 |
| SpCasp | >ACH97121.1 |
| AqCasp | >XP_019854115.1 |
| HvXIAP | >NP_001296622.1 |
| SpXIAP | >XP_022795646.1 |
| AqXIAP | >XP_011406381.1 |
| HsXIAP | >NP_001158.2 |
| AcCaspA | >PP704497 |
| AcCaspB | >PP704498 |
| AcCCaspA | >PP704499 |
| AcCCaspB | >PP704500 |
| AcXIAP | >PP704501 |

**S2 Table**

| Primer | Sequence (5’-3’) | Target gene |
|---|---|---|
| AcCaspA-F | ATGGCTTCCAGAGGAAAGCGT | AcCaspA |
| AcCaspA-R | TTATTGGTCTTTGTAGAAAAA | AcCaspA |
| AcCaspB-F | ATGGCGGATGAGACGGACGCT | AcCaspB |
| AcCaspB-R | TCATAAAAATTTCAACAGTTT | AcCaspB |
| AcCCaspA-F | ATGGTGGGAATGGAAGAAAAG | AcCCaspA |
| AcCCaspA-R | TTAAACACTATTTAAAGGTCC | AcCCaspA |
| AcCCaspB-F | ATGCGCGAGCAAATAAGACAA | AcCCaspB |
| AcCCaspB-R | TCATTCACTGTTTGGGTGAAA | AcCCaspB |
| AcXIAP-F | ATGGCTACAAGTGTGGACTAT | AcXIAP |
| AcXIAP-R | CTACGAAAGATAGATCCTAGA | AcXIAP |
| RT-HvXIAP-F | TGAGCATACTCGGCTGCAAA | HvXIAP |
| RT-HvXIAP-R | CCTATGTTCACCCCAGGCAG | HvXIAP |
| RT-Hvactin-F | CTCTGGTGATGGTGTGTCCC | Hydra actin |
| RT-Hvactin-R | CAGAGCTTGAGGCAGCAGTA | Hydra actin |
